# Analysis the molecular similarity of least common amino acid sites in ACE2 receptor to predict the potential susceptible species for SARS-CoV-2

**DOI:** 10.1101/2023.10.13.562198

**Authors:** YeZhi Hu, Xin Fan, Arivizhivendhan Kannan Villalan, Shuang Zhang, Fekede Regassa Joka, XiaoDong Wu, HaoNing Wang, XiaoLong Wang

## Abstract

This research offers a bioinformatics approach to forecasting both domestic and wild animals’ likelihood of being susceptible to SARS-CoV-2 infection. Genomic sequencing can resolve phylogenetic relationships between the virus and the susceptible host. The genome sequence of SARS-CoV-2 is highly interactive with the specific sequence region of the ACE2 receptor of the host species. We further evaluate this concept to identify the most important SARS-CoV-2 binding amino acid sites in the ACE2 receptor sequence through the common similarity of the last common amino acid sites (LCAS) in known susceptible host species. Therefore, the SARS-CoV-2 viral genomic interacting key amino acid region in the ACE2 receptor sequence of known susceptible human host was summarized and compared with other reported known SARS-CoV-2 susceptible host species. We identified the 10 most significant amino acid sites for interaction with SARS-CoV-2 infection from the ACE2 receptor sequence region based on the LCAS similarity pattern in known sensitive SARS-CoV-2 hosts. The most significant 10 LCAS were further compared with ACE2 receptor sequences of unknown species to evaluate the similarity of the last common amino acid pattern (LCAP). We predicted the probability of SARS-CoV-2 infection risk in unknown species through the LCAS similarity pattern. This method can be used as a screening tool to assess the risk of SARS-CoV-2 infection in domestic and wild animals to prevent outbreaks of infection.

**Graphical abstract:** 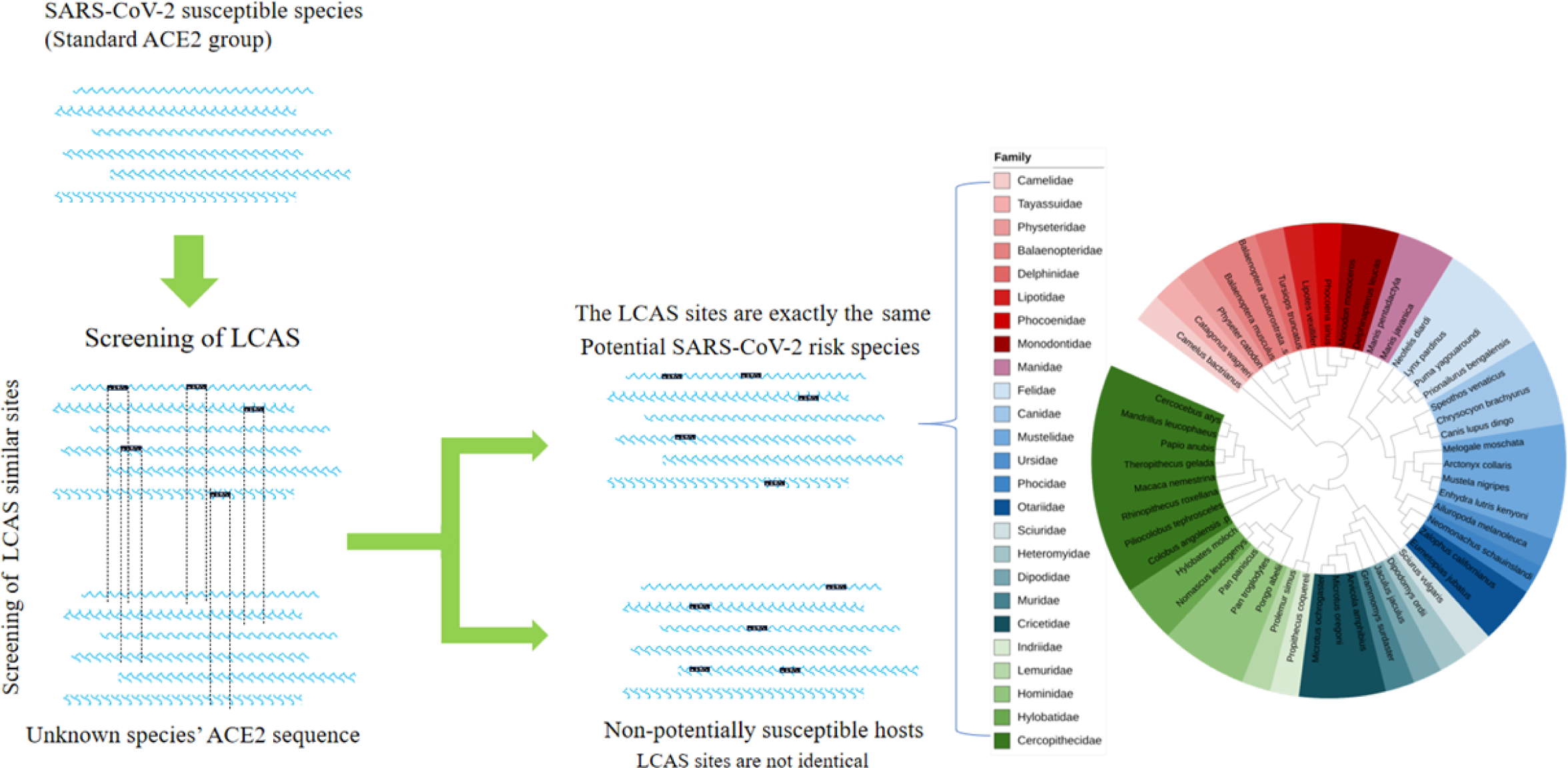

## 1. Introduction

Corona virus disease 2019 (COVID-19) is a highly contagious zoonoses caused by the *Severe Acute Respiratory Syndrome Corona virus 2* (SARS-CoV-2). Since the first case was detected in Wuhan in December 2019, COVID-19 has been spreading globally in a short period of time (1), but the origin of the coronavirus is still unknown. Bats and pangolins have been considered possible natural hosts for SARS-CoV-2, but there is no conclusive evidence (2, 3). The range of SARS-CoV-2 hosts not only humans but also expanding other mammals such as pet cats and minks, which were infected in March and April of 2020 in Belgium and Spain, respectively (4, 5). Subsequently, SARS-CoV-2 infection was detected in ferrets, dogs, golden hamsters, white-tailed deers, rhesus macaques, tigers, lions and so on, as reported by the World Organization for Animal Health (WOAH) (6). An increasing number of mammals are infected with the new coronavirus, indicating the risk of cross-species transmission of SARS-CoV-2. Cross-species transmission of SARS-CoV-2 may lead to further evolution of new hosts and leading to new spread. This poses a serious threat to global public health and biodiversity.

The SARS-CoV-2 viral genome specifically binds to receptors on the surface of host cells, which is a key link in viral infection. So far, the virus has been infecting new species consisting of a specific homologous target receptor capable of binding the SARS-CoV-2 genome. The recognition of SARS-CoV-2 receptors is an important determinant of its transmission between species (7, 8). The specific receptor of the new coronavirus is angiotensin-converting enzyme 2 (ACE2), which is widely expressed in animals as a cell surface receptor. In mammals, ACE2 receptors are distributed in the oral cavity and nasal mucosa, nasopharynx, lung, stomach, small intestine, colon, skin, lymph nodes, thymus, bone marrow, spleen, liver, kidney, and brain (9).

The researcher has extensively studied SARS-CoV-2 in order to determine its host range (10, 11). However, phylogenetic relationships based on whole ACE2 gene comparisons are not a good predictor of mammals at high risk of being infected by SARS-CoV-2 (10, 12). Laboratory infections are not only costly and time-consuming but also have ethical issues, making them difficult to evaluate on a large scale. Therefore, our attention has been turned to the analysis of the key binding domain of ACE2 to SARS-CoV-2. Through extensive research and analysis of the binding structures of SARS-CoV-2 RBD and ACE2, some amino acid residues that are essential for effective binding of ACE2 and SARS-CoV-2 have been identified (8, 13–17).

The analysis of receptor similarity methods is often used to predict the transmission of the virus between species (18). Myeongji Cho’s sequence-based approach suggests that it may be possible to identify virus transmission between hosts without requiring complex structural analysis (12). This method has been used to study the host range of the new coronavirus by predicting the homology of receptor key amino acid sequences, and key binding site methods (10, 11, 19, 20). On this basis, we proposed a new screening approach that involved screening and combining the important Last Common Amino acid Sites (LCAS) in ACE2 from known susceptible hosts, which served as a standard method to evaluate the risk of SARS-CoV-2 infection with unknown species. It can be used as a screening tool and has important scientific implications for discovering potential susceptible hosts of the SARS-CoV-2 virus and assessing its possible transmissibility across species.

## 2. Materials and Methods

### 2.1 SARS-CoV-2 susceptible host collection

Reported SARS-CoV-2 infected species information were collected from the World Organization for Animal Health (WOAH) https://www.woah.org/en/what-we-offer/emergency-preparedness/covid-19/ and literature (6, 21–24). The naturally infected host species and experimentally infected host species information were separately summarized to understand the primary distribution of SARS-CoV-2 infection.

### 2.2 ACE2 receptor sequence collection

The protein sequences of ACE2 from mammalian species were gathered from the National Center for Biotechnology Information (NCBI) Protein Database https://www.ncbi.nlm.nih.gov/ and Uniprot ([UniProt](https://www.uniprot.org/)). The complete ACE2 amino acid sequence of all species were compressed and extracted in FASTA format.

### 2.3 ACE2 receptor data processing

The downloaded sequence file in FASTA format was imported into MAFFT (25) for sequence alignment and duplicate sequences were removed. The human ACE2 receptor sequence was used as a reference to remove gaps and align the sequence in BioEdit (26). The ACE2 receptor data were divided into known susceptible hosts and unknown species for further evaluation. All data were output in FASTA format.

### 2.4 LCAS selection

The key amino acid region of the human ACE2 receptor sequence that strongly binds to SARS-CoV-2 was screened from the literature (7, 8, 10, 13, 14, 27, 28). The standard human amino acid sequence was used to import the important amino acid sections from known susceptible host species into the BioEdit (26) interface. All known susceptible host species were compared to the critical amino acid regions of the standard human receptor sequence. The amino acid sites of known susceptible host species that are commonly similar to human amino acid sequences were separated and named as Last common amino acid sites (LCAS).

### 2.5 Analysis of potentially susceptible hosts

The ACE2 receptor sequence from an unidentified vulnerable host species was matched to the LCAS from recognized susceptible host species. The ACE2 amino acid sequences of known and unknown hosts were extensively analyzed to find the similarity patterns of amino acids. The unknown susceptible species were separated based on a resemblance pattern to the LCAS of known species. Unknown species with entirely identical LCAS patterns were classified as potentially vulnerable hosts; species with variances were designated as non-possibly susceptible hosts.

The MEGA11 software adjacency method (Neighbor Joining Method NJ) was used to construct a phylogenetic tree of potentially susceptible hosts. The average distance of each species in the NJ phylogenic tree was constructed between 0 and 1. We perform a bootstrap test with 1000 replicates to build a phylogenetic tree.

## 3. Result

### 3.1 Collection of SARS-CoV-2 susceptible hosts

The list of animals infected with SARS-CoV-2 was collected from WOAH reports and literature. The results reveal that a total of 55 species were infected with SARS-CoV-2, including 34 species from 14 families that were infected from natural sources (Table 1) and 19 species from 11 families that were infected under experimental conditions. (Table 2) Known susceptibility host statistics (Fig 1).

**Fig 1.**
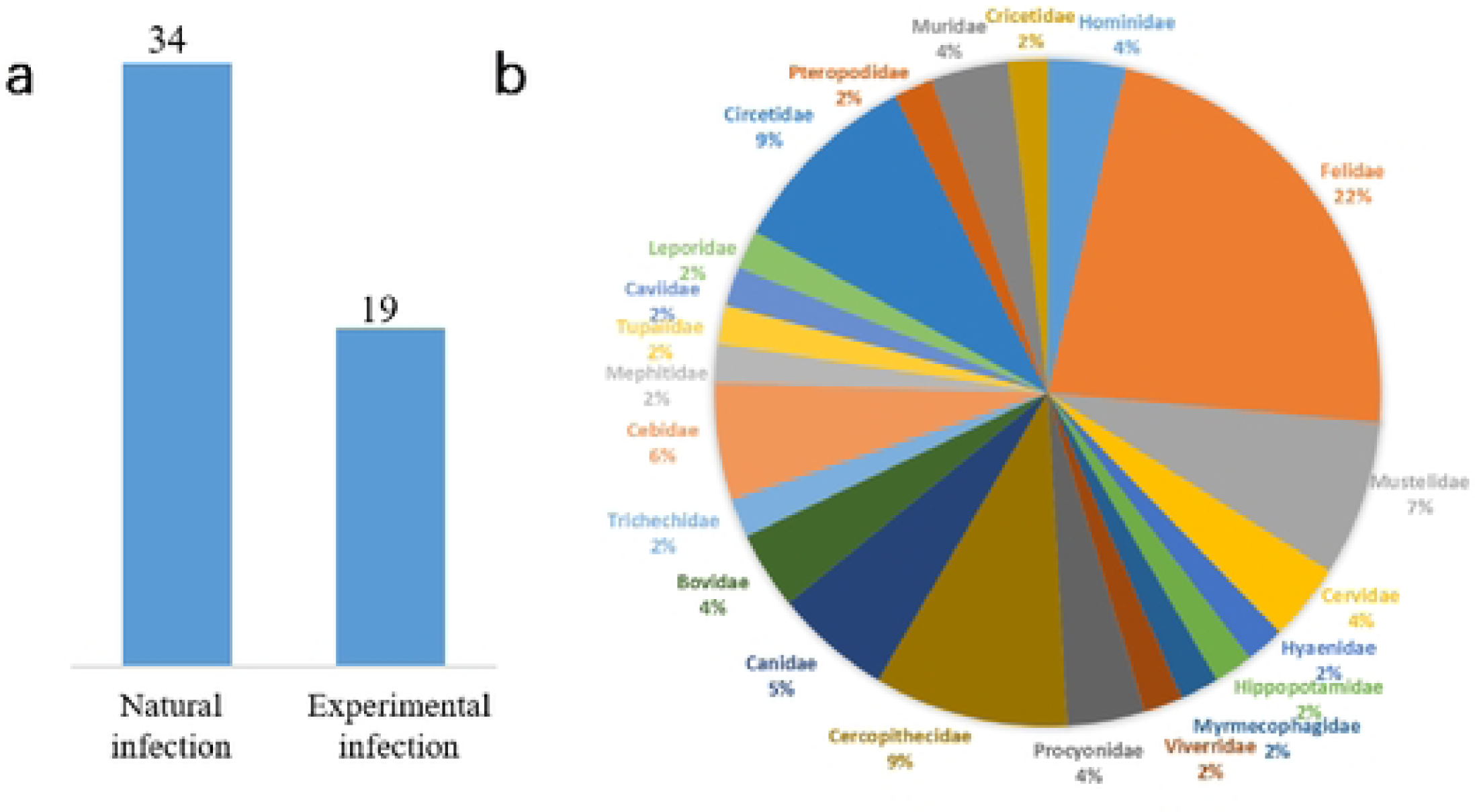
(a) COVID-19 reported species infected by natural and experimental condition and (b) Percentage of animal species in families infected with COVID-19.

**Table 1.**
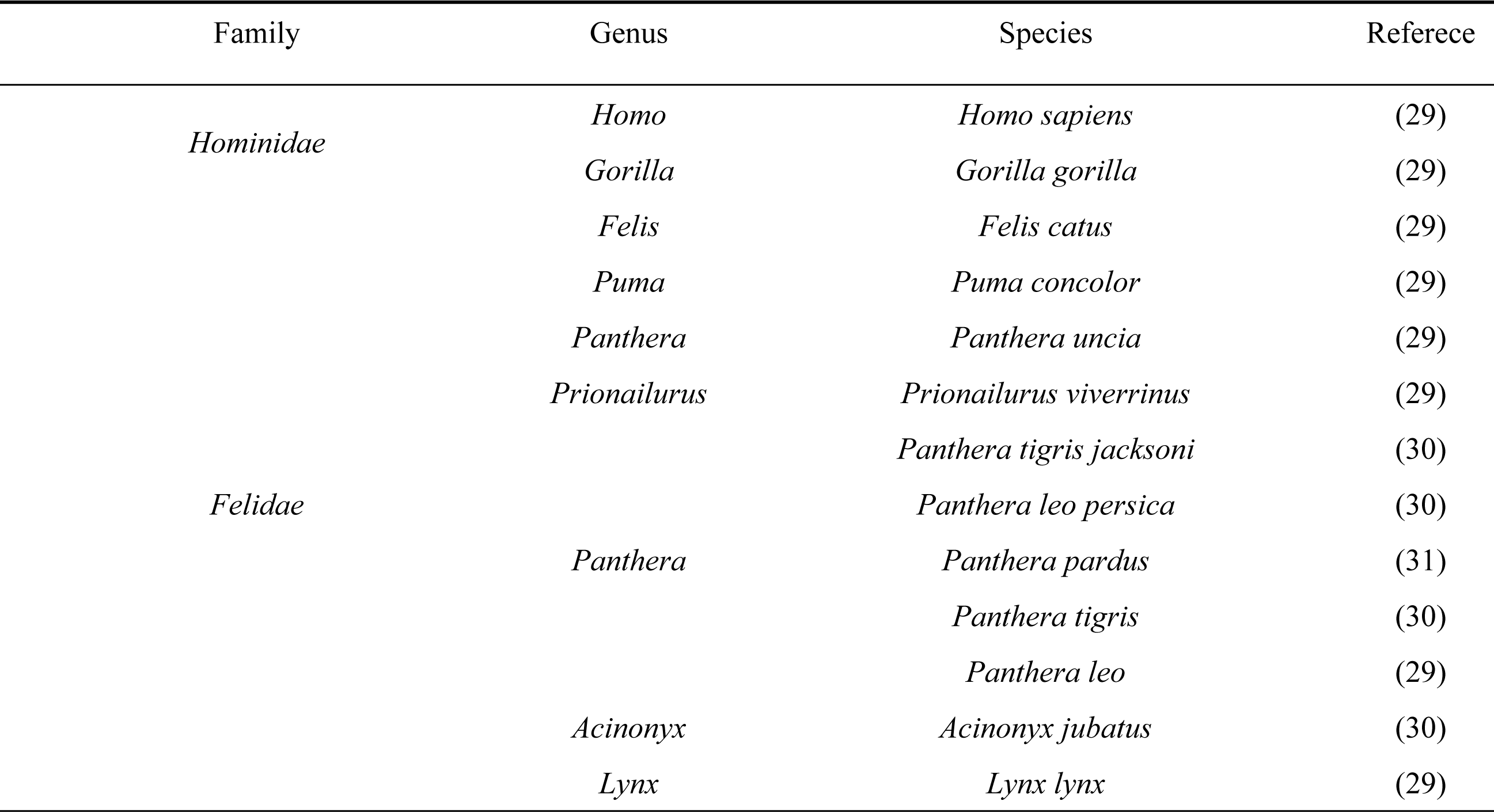
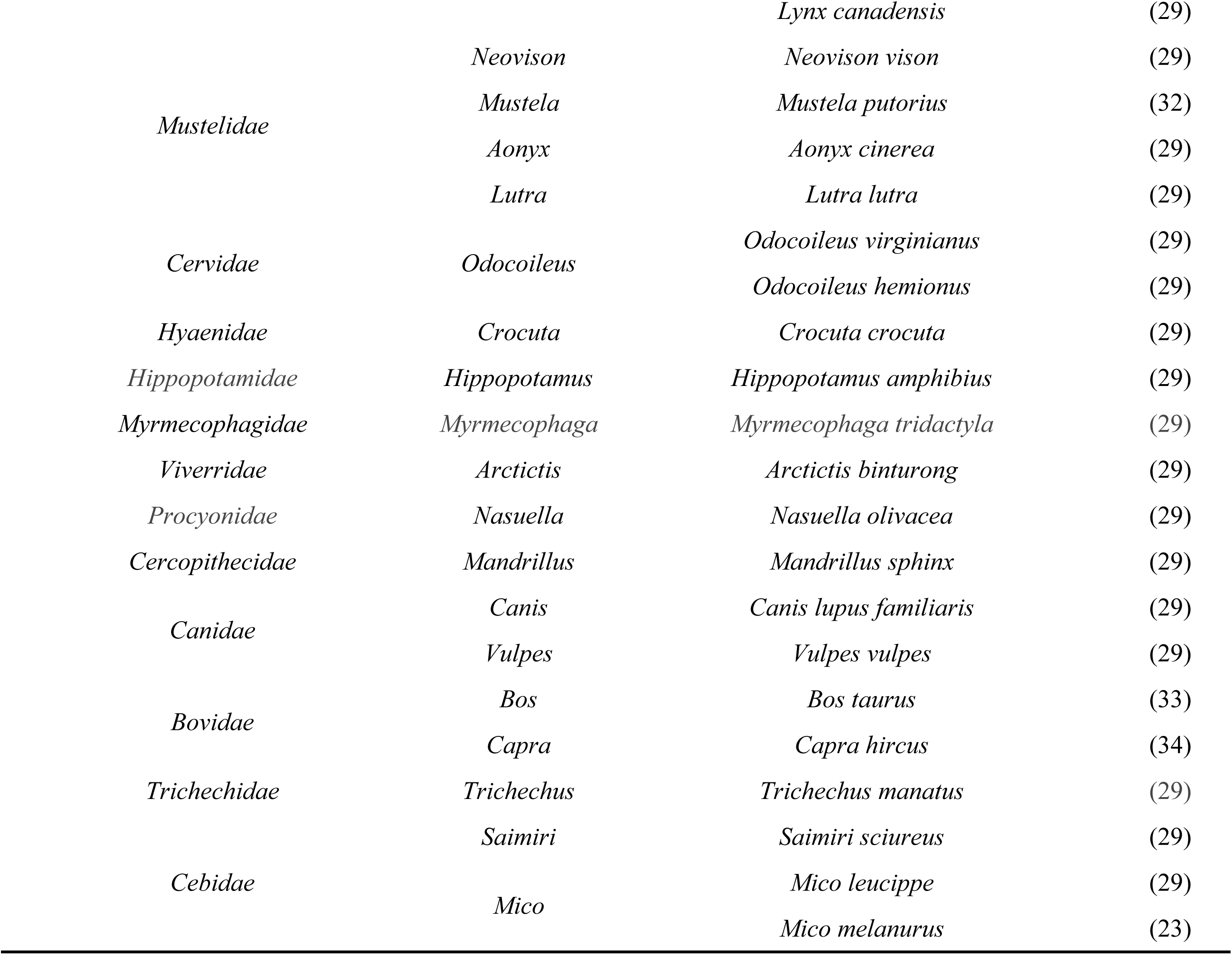
Animals naturally infected with SARS-CoV-2.

**Table 2.**
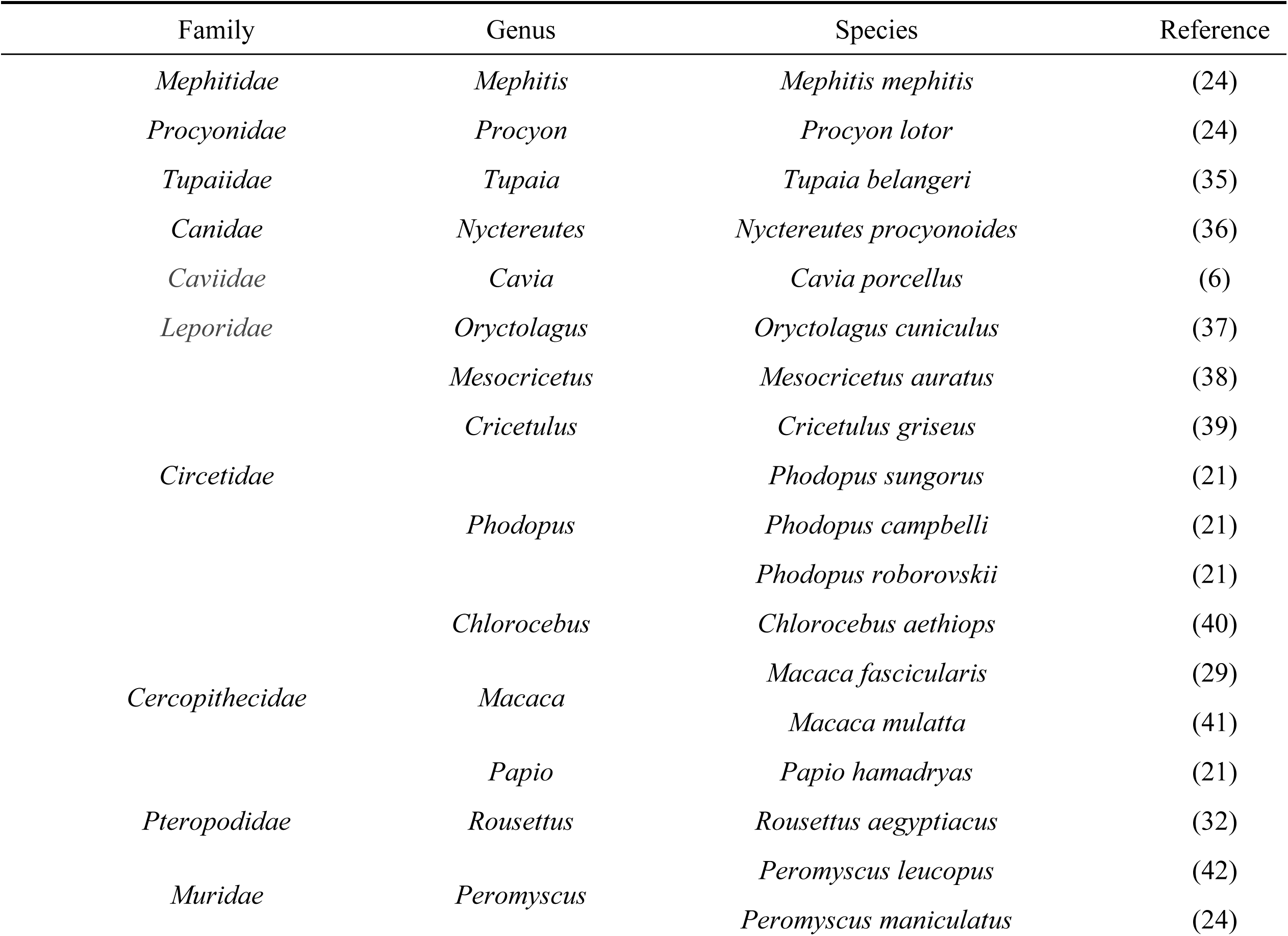
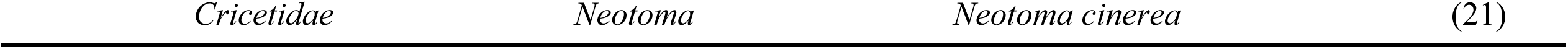
Animals experimentally infected with SARS-CoV-2.

### 3.2 Collection of the ACE2 receptor sequence

We collected 407 ACE2 protein receptor sequences from various species from the Uniprot database. We scrutinized 86 complete ACE2 protein sequences after eliminating incomplete and duplicate sequences. In addition, we obtained 23 complete ACE2 protein sequences from the NCBI database. Finally, 109 ACE2 protein sequences from 45 families were selected for further evaluation to predict the potential risk host for SARS-CoV-2 infection (Table 3).

**Table 3.**
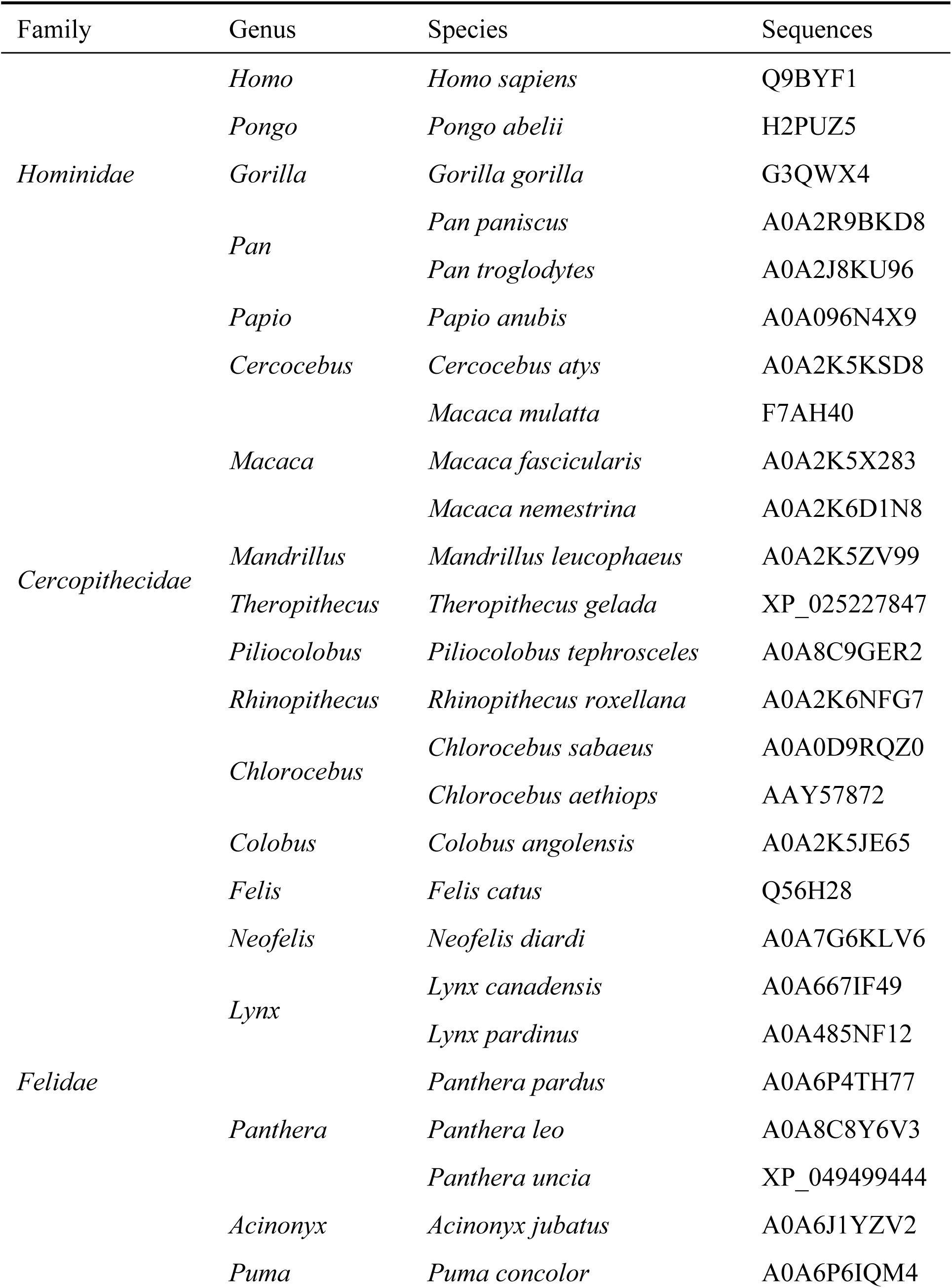
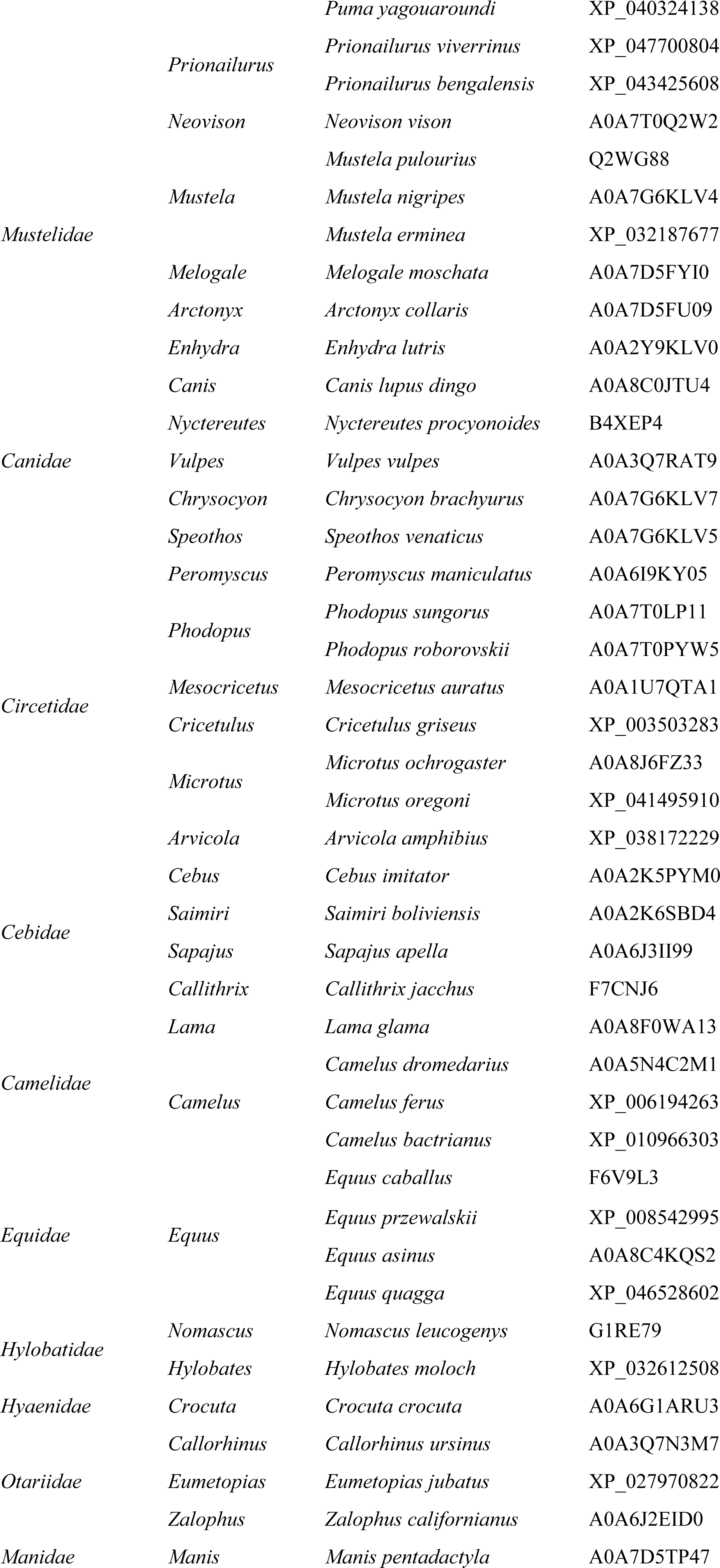
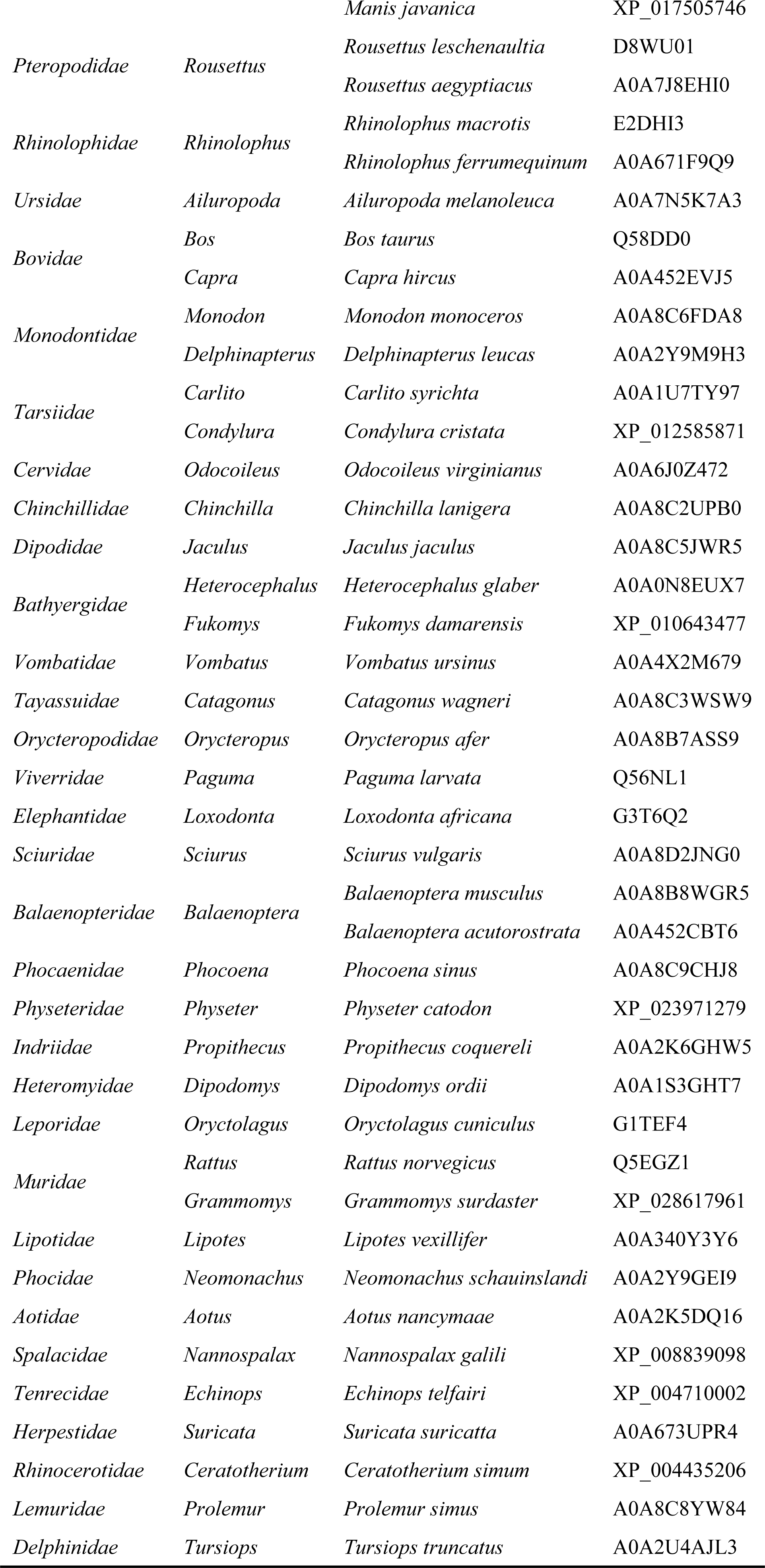
List of ACE2 receptor sequences species used for prediction.

### 3.3 Processing of ACE2 receptor data

We classified 109 ACE2 receptor sequences by dividing them into two groups: the known vulnerable hosts group (29 species in 10 families) and the unknown susceptible hosts group (80 species in 35 families) (Tables 1 and 2). We screened 29 species of ACE2 receptor sequences from 109 as known to be sensitive to SARS-CoV-2. The key regions of the ACE2 receptor sequence in the human ACE2 receptor have been selected for further study (Table 4).

**Table 4.**
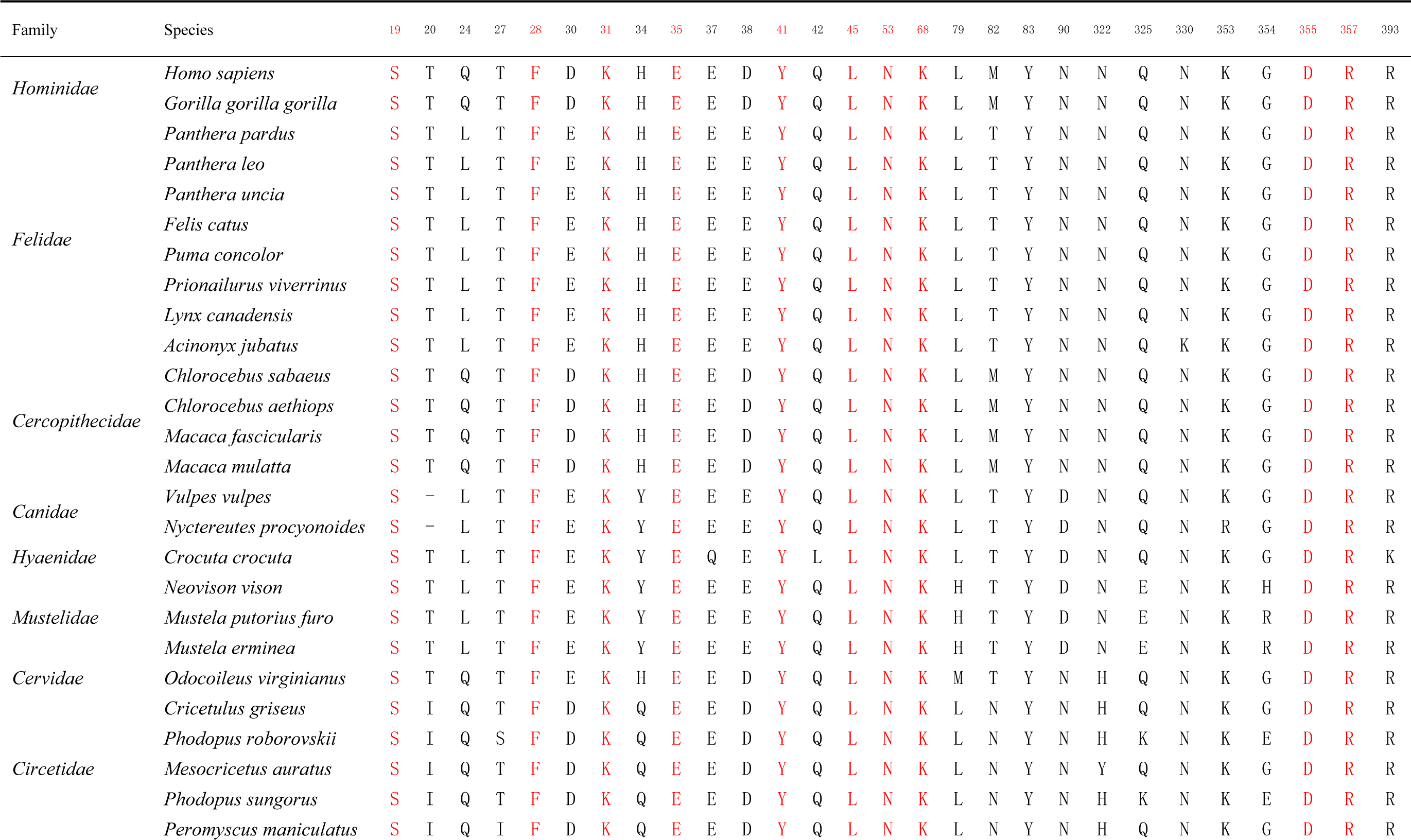
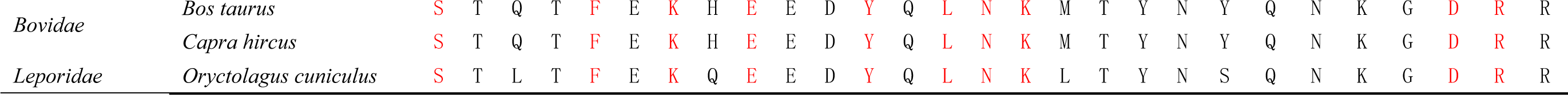
Analysis the similarity of LCAS in conserved loci of known susceptible hosts.

### 3.4 Screening of LCAP

The key regions of the ACE2 receptor sequence in the human ACE2 receptor was compared to the known susceptible to SARS-CoV-2 (Table 4). As a result of the comparison, the 10 most common amino acid sites such as 19, 28, 31, 35, 41, 45, 53, 68, 355 and 357 were identified and used them to further screen the potential risk host for SARS-CoV-2 (Fig 2).

**Fig2.**
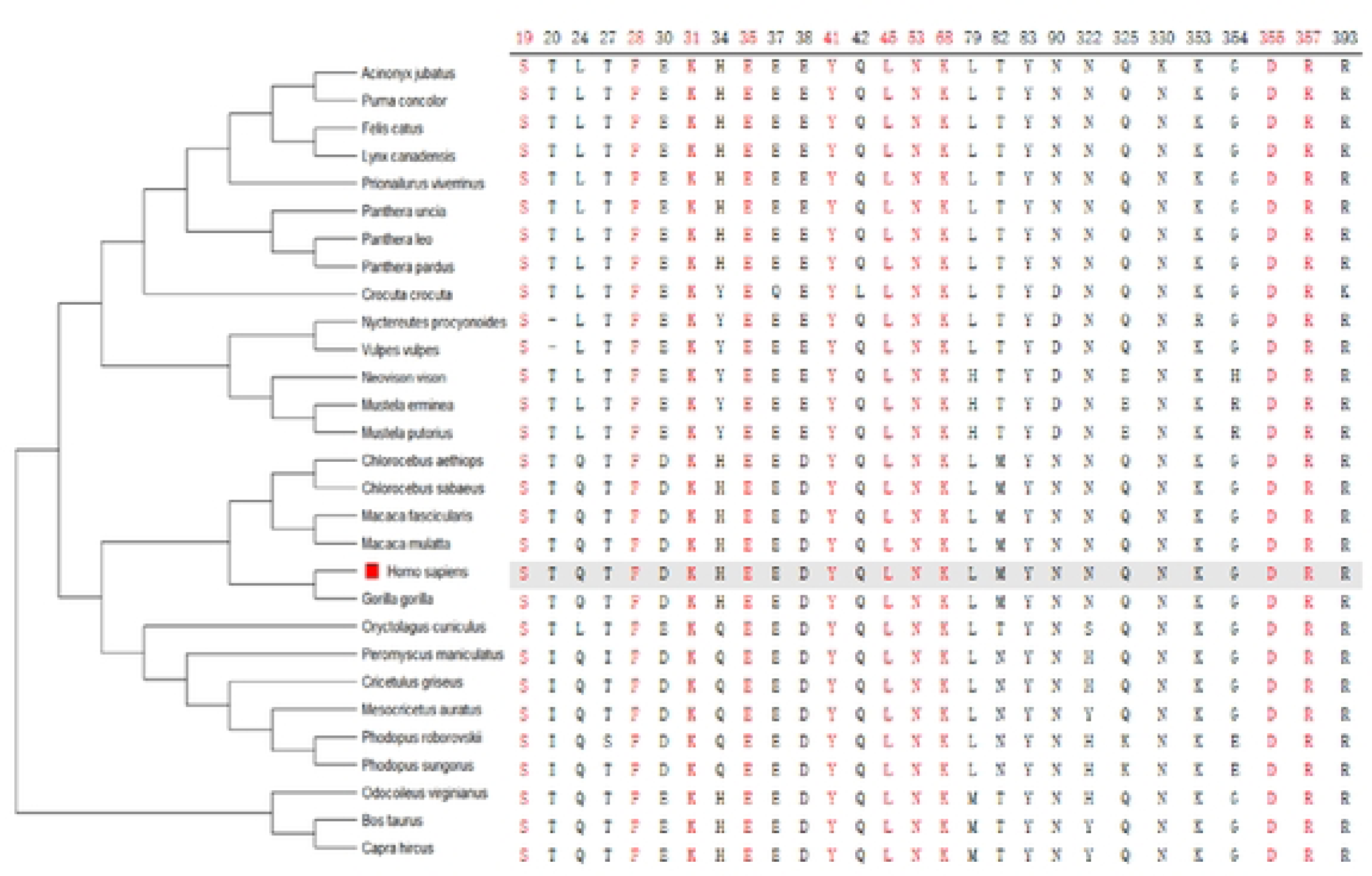
The key structural domains in ACE2 from known SARS-CoV-2 susceptible species.

### 3.5 Analysis of potentially susceptible hosts

In this study, ACE sequences from 80 unknown species were compared to 10 LCAS and their similarity pattern was examined. The ACE2 receptor sequences of 49 non-susceptible species in 25 families were completely similar to those of 10 LCAS of known sensitive species, and they were deemed possible susceptible hosts (Table 5). 31 species from 21 families were considered non-potential susceptible hosts because they were not related to the 10 LCAS. Potential susceptible hosts are located mostly in the orders Primates, Carnivora, Rodentia, and Artiodactyla, which indicates that closely related animals are more likely to be infected with the novel coronavirus. It illustrates the evolutionary links between potentially susceptible risk hosts (Fig 3).

**Fig 3.**
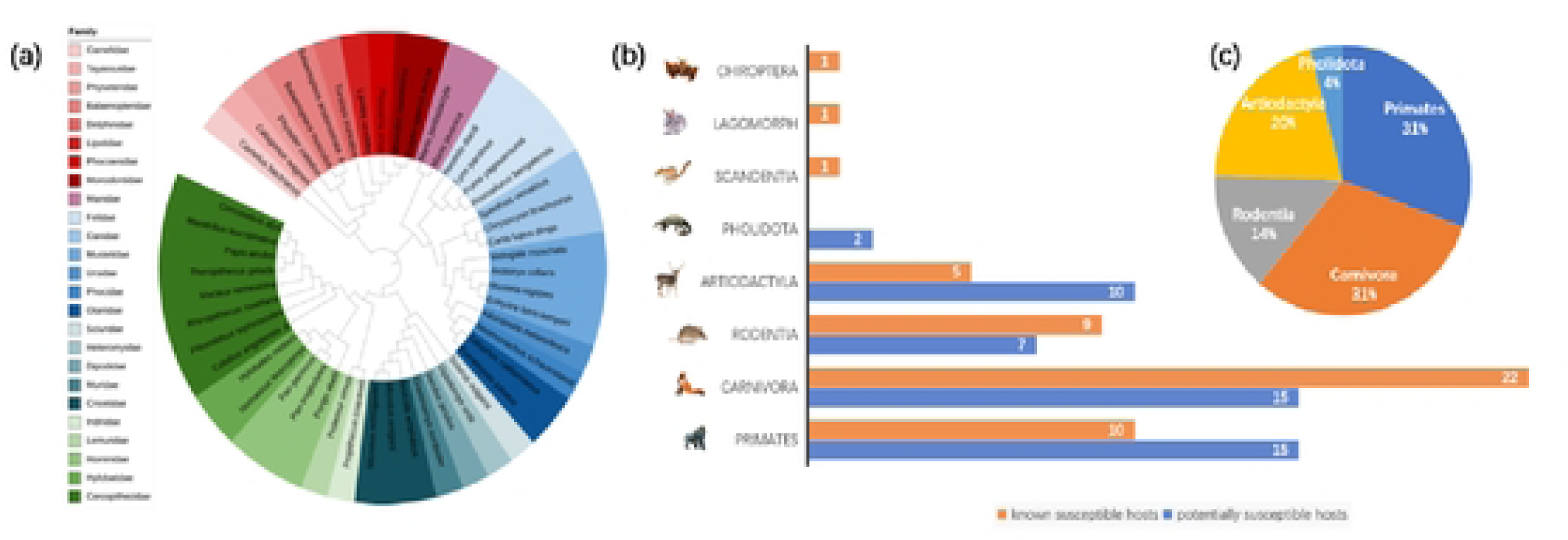
(a) The MEGA11 module calculates the IQ-TREE optimal model to build a phylogenetic tree. iTOL shows the percentage of the total number of species in the outer circle by order, including proportion, and the number of species in the inner circle by family. (b) Shows the number of species in each order in a two-dimensional bar chart. (c) Percentage of animal species in the classification orders potential risk for COVID 19.

**Table 5.**
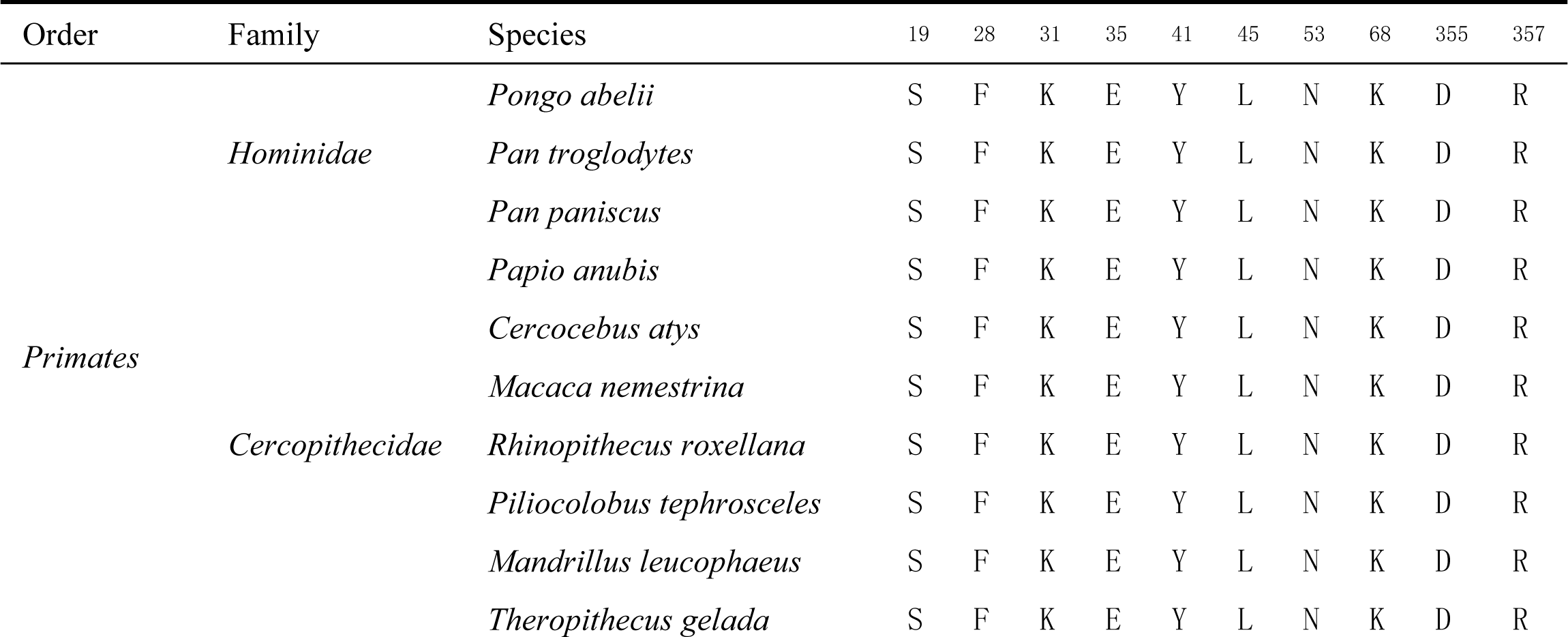
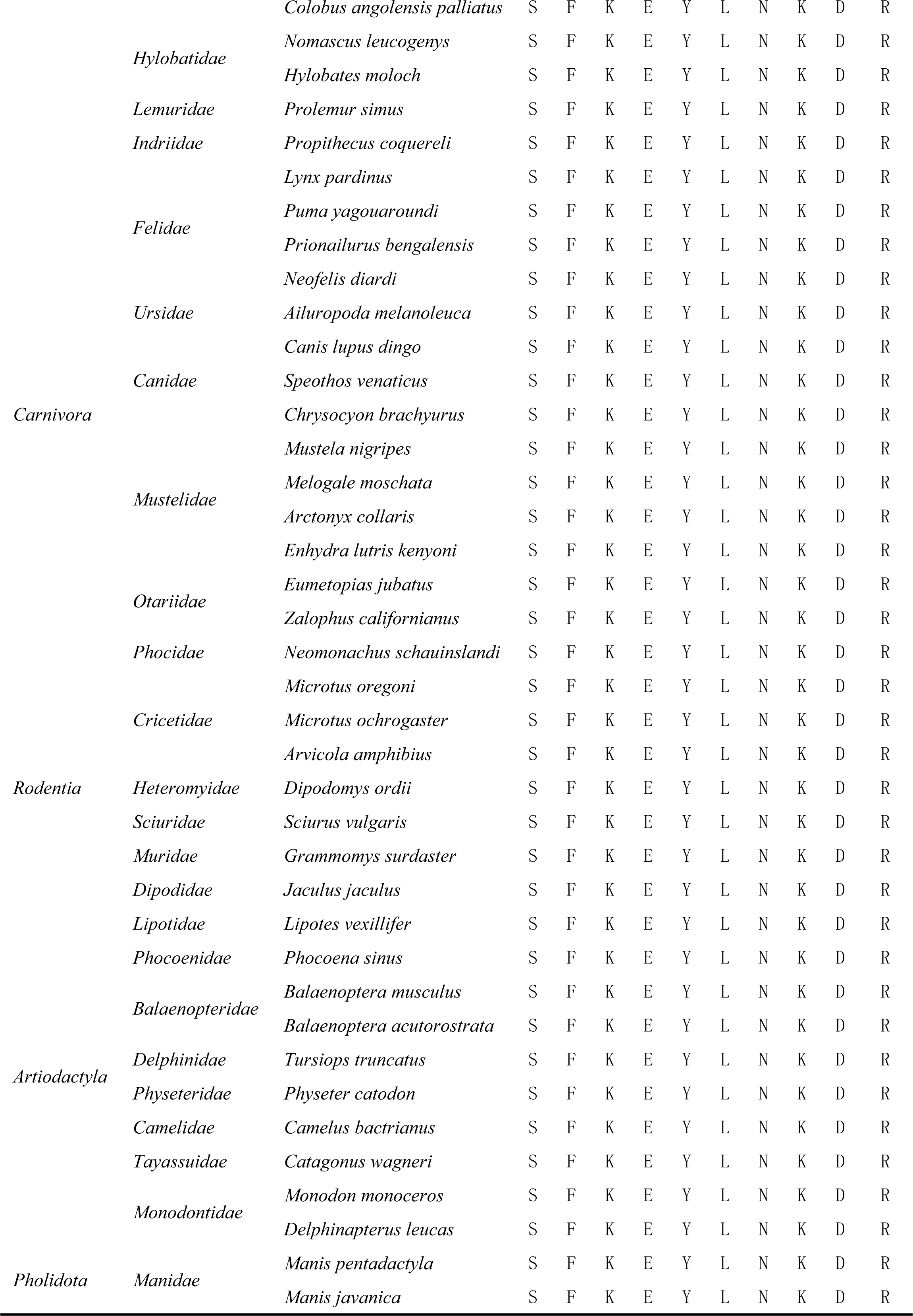
LCAS of potentially susceptible hosts.

## 4. Discussion

In this article, we predicted that the susceptible hosts of SARS-CoV-2 were mainly humanoids, cats, monkeys, and mustelids. Most of these infected animals have been in contact with humans (30). Nevertheless, farmed minks have been shown not only to be susceptible to a natural infection but also to develop severe illness and transmit SARS-CoV-2 to other minks and humans (43). Especially some pet animals have been reported to be infected with the new coronavirus (22), which caused serious concerns because it could be transmitted again to human.

Currently, polymerase chain reaction (PCR) based laboratory testing for SARS-CoV-2 is a standard practice for detecting viral infection and the development of COVID-19 in humans. However, there has been limited research on viral infections, particularly in wildlife animals (20). The increasing number of wild and domestic animals infected with SARS-CoV-2 is a challenge for the rethinking of outbreak control strategies in the post-epidemic era and the response to future emerging infectious diseases. We used open-access bioinformatics tools to predict susceptibility to SARS-CoV-2 infection in domestic and wild animals. For this purpose, we performed a comparative analysis of the ACE2 receptor-specific protein sequences of 109 species. We identified important 10 key amino acid sites that are commonly located in known SARS-CoV-2 susceptible species as reference standards for the analysis and used them to identify the potential risk host. The results reveal that 49 species were potentially susceptible hosts, and 31 species were non-susceptible hosts. Most of the potential susceptible hosts are distributed in the same order as the known susceptible hosts, indicating to some extent that closely related species are more susceptible. Two target species (*Manis pentadactyla* and *Manis javanica*) in the prediction results have not been reported so far, indicating that while focusing on closely related species, it is necessary to pay attention to other target species and protect animals on a larger scale.

The results show that not all primates are susceptible to SARS-CoV-2. The 41-position key amino acids of ACE2 receptors in primates, *capuchinidae*, night monkeys and marmosets are different from humans. A large number of studies have confirmed that 41-position amino acid mutations may break key hydrogen bonds, reducing the binding capacity of SARS-CoV-2 to ACE2 (12, 44); Bats are generally considered to be the main natural hosts of the new coronavirus, but the 35 amino acids of *Rhinolophus macrotis* and *Rhinolophus ferrumequinum* of the *Rhinolophidae* family are different from humans (28). The mutations in E35K can reduce the binding capacity of SARS-CoV-2. Jun Lan et. al., (2021) found that ACE2 of *Rhinolophus ferrumequinum* cannot mediate the entry of the new coronavirus (45). It suggests that not all bats are susceptible to the new corona virus. Assessing the susceptibility of various bat species to the new coronavirus is the first step in the traceability process for bats, which can significantly reduce the challenges in tracing the new coronavirus. Equestrians are non-susceptible species and may be associated with differences in amino acids 41 and 68, which have been shown to reduce SARS-CoV-2 binding to receptors (28).

The primary structure of the receptor protein influences the host’s susceptibility to cross-species viruses, making it possible to identify viral transmission between hosts without the need for complex structural analysis (46). Key receptor amino acids have been used in several kinds of studies to predict susceptible species. Yinghui Liu et. al., (2020) predicted 26 mammals based on 20 key amino acid sites, and analyzed the electron microstructure of hACE2 bound to SARS-CoV-2 RBD to verify the prediction results (11). Eighty mammals were shown to be susceptible to SARS-CoV-2 through an investigation of the conservativity of five residues in the two viral binding sites of the ACE2 receptor (12). Yulong Wei et. al., (2021) selected 15 key amino acid sites to analyze the similarity of ACE receptor amino acid sequences in 131 mammals (10).

In this study, a minimum number of key amino acid loci were selected based on the LCAS of known susceptible hosts, which greatly reduces the complexity of the work and allows for rapid and more accurate prediction of potentially susceptible hosts for the new coronavirus. Genetic variations in the host receptor ACE2 may also contribute to susceptibility or resistance against the viral infection, depending on how the variations in spike protein influence the cross-species transmission of the virus. Studies have proved that after genetic mutations in S19, K31, E35, Y41, K68, and D355, the binding capacity of the virus to the receptor decreases (27, 28). The predicted results are almost consistent with the results of other studies (19), indicating the accuracy of the results. The predicted results are almost consistent with the results of other studies (2), indicating the accuracy of the results. This method is simple and accurate, which can provide ideas to predicting the potential susceptible hosts in the early stages of disease outbreaks. It supports protective preventive measures for potential hosts in advance to control future outbreaks and reduce animal infections. The constant mutation of coronavirus increases its ability to bind to the ACE2 receptor as well as resist the immune response (47). For example, N501Y can form a new interaction with the ACE2 receptor Y41, and it is widely present in mutants (48). Especially the mutated Omicron strain S residue Y501 stacking interaction with the T-shaped π–π of Y41 in the ACE2 residue. The Q493R and Q498R mutations introduce two new salt bridges, such as E35 and E38, respectively replacing hydrogen bond formation and remodelling the electrostatic interactions with the ACE2 receptor of Wuhan-Hu-1 RBD. S477N leads to the formation of new hydrogen bonds between the asparagine side chain and the ACE2 S19 backbone amine and carbonyl groups (47, 49, 50). These interactions illustrate that key amino acid sites on the ACE2 receptor are important for viral binding.

In this study, we only considered key amino acid sites of virus-receptor interactions to predict susceptibility. However, the viral entry into host cells and replication were influenced by many other factors, such as cathepsin TMPRSS2 or CTSL1, and ADAM-17 (51). Therefore, key amino acid sites alone are not sufficient. Laboratory validation and clinical observation of the listed animals are necessary to confirm their susceptibility to SARS-CoV-2 infection.

## Acknowledgements

The authors thank everyone who has participated in the data collection for this study.

## Notes

### Competing Interest Statement

The authors have declared no competing interest.

